# Evidence for a role of calcium in STING signaling

**DOI:** 10.1101/145854

**Authors:** Sujeong Kim, Peter Koch, Lingyin Li, Leonid Peshkin, Timothy J. Mitchison

**Affiliations:** Department of Systems Biology, Harvard Medical School, Boston, MA 02115, USA.

## Abstract

STING, an ER resident cyclic dinucleotide (CDN) receptor, plays an important role in innate immune response signaling. Upon binding to CDNs, it activates the TBK1-IRF3 signaling axis, which stimulates gene expression including interferon beta. We hypothesized that the ER localization of STING reflects a role for calcium mobilization in its signaling. To test this hypothesis, we treated mouse cells with two STING agonists, the synthetic drug DMXAA and a natural ligand, cyclic GMP-AMP (cGAMP), and measured intracellular calcium. Both triggered a rapid rise in intracellular calcium that was partially inhibited by STING depletion. Intracellular calcium chelation blocked DMXAA induced signaling downstream of STING activation, but had no effect on cGAMP induced signaling. We propose that intracellular calcium plays an important role in the response of the STING pathway. In response to DMXAA, calcium is mobilized form the ER and required for signaling. In the case of cGAMP calcium is mobilized but not required. This difference could be explained by alternative modes of STING activation for the two ligands, or a combination of STING-dependent and -independent actions of extracellular cGAMP.

## Introduction

Cyclic dinucleotides (CDNs) are synthesized by bacteria, and constitute a form of Pathogen-Associated Molecular Patterns (PAMP) recognized by the innate immune system. Cytosolic DNA, a marker of viral infection, triggers production of a related CDN, 2’3’-cGAMP (phosphodiester linkages between 2’-OH of GMP and 5’-phosphate of AMP, and 3’-OH of AMP and 5’-phosphate of GMP) by the infected cell through activation of the enzyme cyclic GTP ATP synthase (cGAS) [1,2]. The best characterized CDN receptor is STING (stimulator of IFN genes), an endoplasmic reticulum (ER) resident protein with four membrane-spanning helices [3,4]. STING, which is dimeric, binds a single CDN in a cytosolic ligand-binding domain [4]. CDN binding results in conformational changes of STING and recruitment of TANK binding kinase 1 (TBK1) and interferon regulatory factor 3 (IRF3)[5]. TBK1 phosphorylates IRF3 on S396, causing nuclear translocation and transcriptional activation of many genes including interferon beta [6,7]. Precisely how STING activates transcription is still poorly understood. Recent reports showed that STING has multiple phosphorylation sites involved in its activation. Translocation of STING from ER to golgi and STING degradation that is regulated by ubiquitin E3 ligases have also been implicated in regulating the STING pathway [8,9,10].

One unsolved puzzle is why STING is an ER transmembrane protein – does the ER membrane simply provide a physical environment for STING activation or does it play other roles? In many signaling pathways, the ER serves as a source of calcium ions, which are released into the cytoplasm by IP3 receptors [11]. Given its location in the ER, we hypothesized that STING might also use calcium release from the ER as part of its signaling mechanism.

Calcium plays many roles in signaling during innate and adaptive immunity. For example, an increase in cytosolic calcium is often part of activating immune cells to execute some effector function or initiate proliferation [12,13]. Lipopolysaccharides (LPS) have been shown to induce the production of diacylglycerol (DAG) in TLR4 dependent manners. This leads to increased intracellular calcium that is implicated in control of gene expression downstream of TLR4 activation [14,15]. Recently, crystal structure of STING has been published by Li group and they showed that STING has two calcium ion binding sites at the loop structure [16]. However, a possible role of calcium in the STING pathway has not, to our knowledge, been investigated. Schindler group also showed that CDNs added to the outside of cells can activate inflammasomes by a mechanisms dependent on potassium and calcium fluxes, but in this case the pathway was independent of STING [17].

To test if STING activation triggers calcium transients, we needed ligands that could be added to the outside of cells and that rapidly activate STING. STING is typically activated by transfecting cells with DNA, but this takes minutes to hours, and the transfection reagents themselves would likely trigger calcium transients through indirect mechanisms. Several small molecule drugs are agonist ligands for mouse, but not human, STING. These include the anti-cancer flavonoids Flavone acetic acid (FAA) and DMXAA [18,19,28], and the anti-viral CMA (10-carboxymethyl-9-acridanone) [20]. On the basis of binding studies, crystal structures and pathway analysis, these compounds are thought to activate mouse STING in the same way as the natural cyclic dinucleotide ligands [18,21]. In addition, CDNs themselves are surprisingly active when added to the outside of cells. They cause rapid activation of the STING pathway [21] as well as STING-independent effects leading to inflammasome activation [17]. We used both ligand classes to test for a role of calcium transients in STING signaling.

## Materials and Methods Cell culture and reagents

Raw264.7 and L929 cells were grown in DMEM (Cellgro) supplemented with 10% FBS (GIBCO) (v/v) and 1% of penicillin-streptomycin (Cellgro). DMXAA and XAA-8Me were prepared as previously described [18]. Cyclic di-nucleotides cyclase construct (DncV), cloned out from cholera ToxT was generous gift from Dr. Mekalanos [22]. 3’3’-Cyclic GMP-AMP (phosphodiester linkages between 3’- OH of GMP and 5’-phosphate of AMP, and 3’-OH of AMP and 5’-phosphate of GMP) was synthesized as previously reported [22] and directly added to the medium for treatment. Rabbit polyclonal antibodies against TBK, pTBK (S172), pIRF3 (S396), were purchased from Cell Signaling Technology. STING rabbit polyclonal antibody was raised against the C-terminal domain of STING (139-378) by Cocalico Biologicals, Inc. BAPTA-AM was purchased from Invitrogen and dissolved in DMSO. 2-APB and Xestospingin C (Cayman Chemical) were also dissolved in DMSO.

### Calcium imaging and imaging analysis

We obtained the genetic calcium reporter, gCaMP3-GFP from Addgene. Raw264.7 and L929 cells were transfected with the gCaMP3-GFP plasmid and selected for more than one month. We seeded cells on to fibronectin coated glass bottom 35mm petri dishes (MatTek) a day before imaging. A few hours prior to imaging, medium was replaced with phenol-red free DMEM supplemented with 10% (v/v) FBS, 1% (v/v) antibiotics and 25mM HEPES (pH7.2). All images were collected with a Nikon Ti automated inverted microscope with perfect focus and a CO_2_ and humidity controlled 37°C incubation chamber.

For image analysis we analyzed fluorescence intensities of individual cells over time with Image J and Matlab.

### Centrifugal ultrafiltration assay

Mouse STING (amino acids 139-378), purified as previously described[18], was diluted to a concentration of 200 μM with Tris buffer (20 mM Tris, 150 mM NaCl, pH 7.5). DMXAA and cGAMP were diluted to the same concentration as the protein in Tris-buffer. Equal volumes (20 μL) of small molecules and STING protein were mixed and loaded onto centrifugal filtration devices (NANOSEP 3K OMEGA). Absorbance measurements from the flow-through of a one minute spin were read on a NanoDrop spectrophotometer (ND-1000). Unbound DMXAA and cGAMP were monitored by their UV absorbance at 245 nm and 254 nm respectively.

### Immuno-staining

We fixed cells using 4% paraformaldehyde for 20 min at room temperature. After permeabilization, cells were incubated with anti-STING antibody overnight and visualized with a FITC labeled secondary antibody. DAPI was used to visualize the nucleus. Images were taken by spinning disk confocal microscopy.

### IP1 assay

IP-one assay kit was purchased from Cisbio Assays. In brief, cells were seeded in a 96-well white plate (Greiner Bio-one). The next day, cells were treated with DMXAA or UDP (Sigma) in the presence of LiCl. After cell lysis, IP1 levels were measured by the competition of endogenous IP1 and deuterium labeled IP1. FRET signal was measured by EnVision 3 (Perkin Elmer).

## Results

### STING agonists induce transient calcium responses

To investigate the role of calcium in STING signaling we expressed the calcium reporter gCaMP3-GFP [23] in the Raw264.7 mouse macrophage cell line that is commonly used to study STING signaling.

This fluorescence intensity of this reporter increases upon calcium binding, but with no spectral changes; its K_D_ for binding calcium ions is 405 ± 9nM [23]. Figure 1A shows averaged pixel intensity of a single cell over time measured from time-lapse movies of fields of cells taken with a 20x objective. Drug was added at t = 0 and images were taken every 20 seconds for 30 minutes. Raw264.7 cells are macrophage-like and initiate calcium transients when they move or phagocytose [24,25]. This likely explains fluctuations in calcium signal, including spikes, observed under basal conditions (blue line in Fig. 1A). 62% of cells showed this type of calcium pulsing without any treatment. Addition of DMXAA to the medium triggered a large spike in intracellular calcium levels that lasted for a few min (red line in Fig. 1A). To test whether this initial increase was statistically meaningful, we normalized the fluorescent intensities of each cell by its median value. Then we calculated the median intensities of the whole cell population across time (Fig. 1B). The calcium increase immediately following DMXAA addition was clearly apparent in the averaged trace (red line in Fig. 1B). Analyzing across >90 cells, DMXAA caused both an early strong calcium increase, and more subtle longer-term changes (Fig. 1C). DMXAA treated cells pulsed more frequently as judged by both the mean number of pulses (5.9 vs 5.2) and the percentage of cells with more than 3 pulses (67.3% vs 53.3%). We used Kolmogorov-Smirnov test (K-S test) to get *p*-values (*p*-values<0.005).

**Figure 1.**
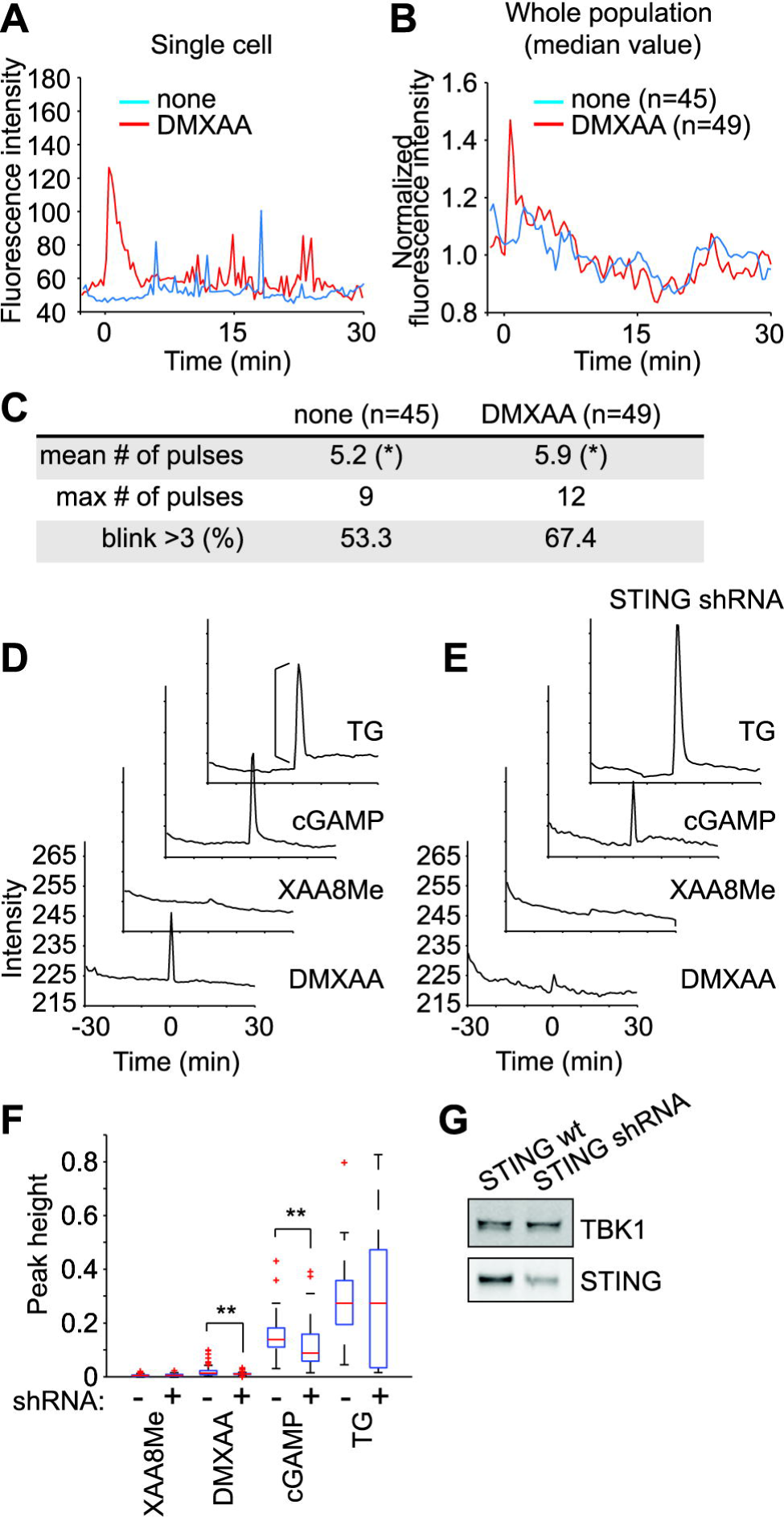
DMXAA and cGAMP increased intracellular calcium dynamics. (A) Representative gCaMP3-GFP intensity profiles with or without DMXAA (250μM) treatment in Raw264.7 cells. (B) Mean intensity profile of the indicated number of cells. (C) Analyzed number of pulses in a single cell for given time in each condition. An asterisk (*) indicated *p*-values< 0.005 using K-S test. (D) Intracellular calcium traces of gCaMP3-GFP expressing L929 cells with (E) or without (D) STING depletion by lenti-virus delivered shRNA. Drugs were added at time 0 at the concentration of DMXAA (250μM), XAA8Me (250μM), cGAMP (50μM) and TG (1μM). The bracket indicated peak height. (F) Peak height analysis. These were sum of two independent experiments. Analyzed number of cells is as followed. In no shRNA treated cells, 101 (XAA8Me), 69 (DMXAA), 44 (cGAMP) and 47 (TG). In STING shRNA treated cells, 103 (XAA8Me), 99 (DMXAA), 77 (cGAMP) and 51 (TG). The intensity before and after adding drug was subtracted to give the peak height as indicated in figure 1D. Double asterisks (**) indicate p<0.0005(K-S test) (G) Western blot analysis showing STING depletion levels in cells in (E).

Next, we tested whether the DMXAA induced calcium increase was mediated by STING. To address this question, we used gCaMP3 expressing L929 cells. This mouse fibrosarcoma cell line has a functional STING signaling pathway and is easier to manipulate than Raw264.7 cells [5]. Images were taken every 30 sec for 30 min before and after the drug treatment. Drugs were added at t = 0. In general, the calcium traces were less noisy in L929 cells. The SERCA (sarco/endoplasmic reticulum Ca^2+^- ATPase) inhibitor, thapsigargin (TG), was used as a positive control for calcium mobilization (Fig. 1D). We observed a strong pulse of intracellular calcium seconds after DMXAA addition, but no subsequent pulses (Fig. 1D). XAA8Me, a structurally similar, inactive analog of DMXAA [26], did not provoke calcium pulses, which argues against some non-specific effect. cGAMP, added to the outside of cells, rapidly induced calcium pulses that were similar in duration to those induced by DMXAA, but tended to have much larger peak heights.

To test if STING was required for calcium pulses induced by DMXAA and cGAMP, we reduced STING expression using lenti-virus delivered shRNA as previously described [18]. By immuno-blot, we estimate STING levels were reduced by ~50% (Fig. 1G). To quantify the calcium signal, we analyzed peak heights (Fig. 1D, 1E). The fluorescence intensity of an individual cell was normalized by its initial intensity value and the maximum intensity was subtracted by the intensity before drug treatment. The peak heights in each condition were plotted as box plot using MATLAB. As MATLAB function description, edges of box indicated 25% and 75% of the population, the central mark (red line in the box) indicated median value, the whiskers are the most extreme data points that are not considered outliers, and red crosses are outliers (Fig. 1F). With or without STING depletion, TG induced calcium release showed similar peak height as expected. STING depletion caused the median value of peak heights of DMXAA treated cells to decrease by ~15.4%, and cGAMP treated cells to decrease by ~36.3%. We used K-S test to get p-values. These data are hard to interpret in any strictly quantitative sense since STING depletion was incomplete and calcium signaling typically involves complex, non-linear signal processing. However, the data do suggest that STING is partially required for the calcium response to both DMXAA and cGAMP.

### Intracellular calcium is required for DMXAA, but not cGAMP, to activate STING

Next, we questioned whether the calcium response was functionally important for STING induced signal transduction. To address this question, treated L929 cells at t = 0 with EGTA, an extracellular calcium scavenger, or BAPTA-AM, a cell-permeable ester that is rapidly hydrolyzed to the calcium chelator, BAPTA, in the cytosol. We then treated with DMXAA at t = 30min. As measured by gCaMP3-GFP imaging, EGTA pretreatment did not block the increase of intracellular calcium induced by DMXAA, whereas BAPTA-AM completely blocked it (Fig. 2A, 2B). This result indicates that the calcium source was an internal reservoir, such as the ER, and not the extracellular medium.

**Figure 2.**
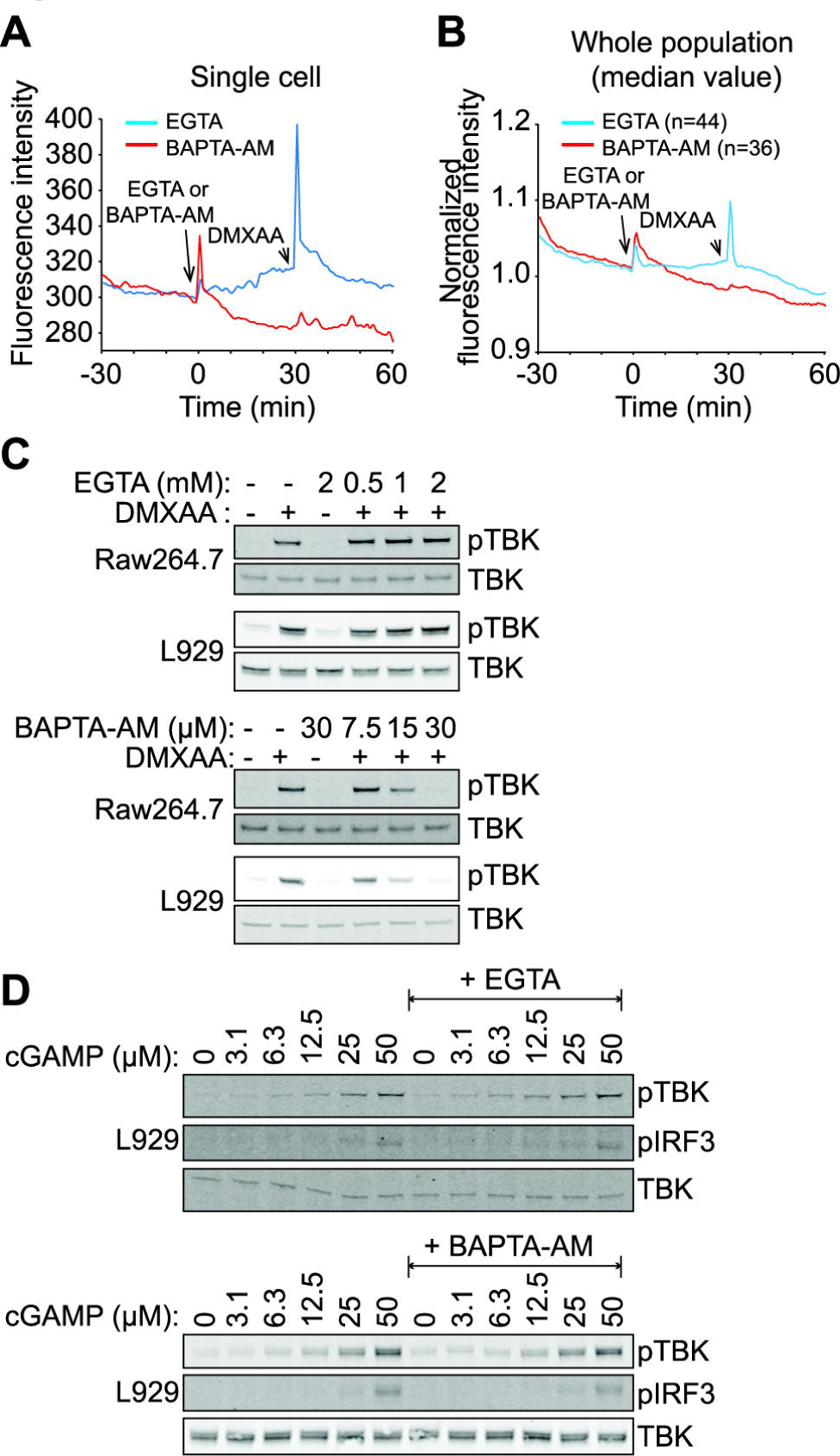
Intracellular calcium inhibition blocked DMXAA induced STING signals but not cGAMP induced signals. (A) Representative calcium trace of gCaMP3-GFP expressing L929 cells. BAPTA-AM (30μM) or EGTA (2mM) was added at the time=0 and followed by DMXAA (250μM) treatment at time=30. (B) Mean intensity profile of indicated number of cells. (C) Cells were pretreated with EGTA or BAPTA-AM for 30 min and followed by DMXAA (250μM) treatment for another 30 min. Phospho-TBK1 and phospho-IRF3 antibodies were used to measure TBK1-IRF3 activation. (D) cGAMP was treated for an hour with or without EGTA (2mM) or BAPTA-AM (30μM) pretreatment for 30 min. TBK1 and IRF3 activity was measured as (A).

To test if calcium increase was important for STING induced signal transduction, we measured TBK1 and IRF3 activation by measuring phosphorylation levels on epitopes previously implicated in pathway activation [5,18] (Fig. 2C). EGTA had no effect, consistent with its lack of effect on calcium levels by imaging. BAPTA-AM completely blocked DMXAA induced TBK1 phosphorylation in dose dependent manners in both Raw264.7 and L929 cell lines (Fig. 2C). Since cGAMP also increased intracellular calcium, we expected BAPTA-AM would also block cGAMP induced STING signaling activation. However, EGTA or BAPTA-AM pretreatment did *not* block cGAMP induced TBK1 or IRF3 phosphorylation (Fig. 2D). These data suggest a rise in intracellular calcium is required for activation of TBK1 and IRF3 by one STING agonist, DMXAA, but not another, extracellular cGAMP.

We next considered the possibility that calcium is differentially required for ligand binding to STING. To address this issue, we added DMXAA or cGAMP to the dimeric ligand-binding domain of mouse STING (139-378), expressed in bacteria and purified, and measured ligand binding with a centrifugal ultrafiltration assay as previously described [18]. When small molecule alone was loaded into a filter with a 3kDa cutoff, unbound small molecules flowed through and were quantified by UV absorbance. Addition of mSTING (139-378) caused partial retention of the ligand above the filter, resulting in a drop in absorbance measured at 245nm for DMXAA and 254nm for cGAMP. Calcium chelation by EGTA had little effect on binding of either DMXAA or cGAMP (Fig. 3A).

**Figure 3.**
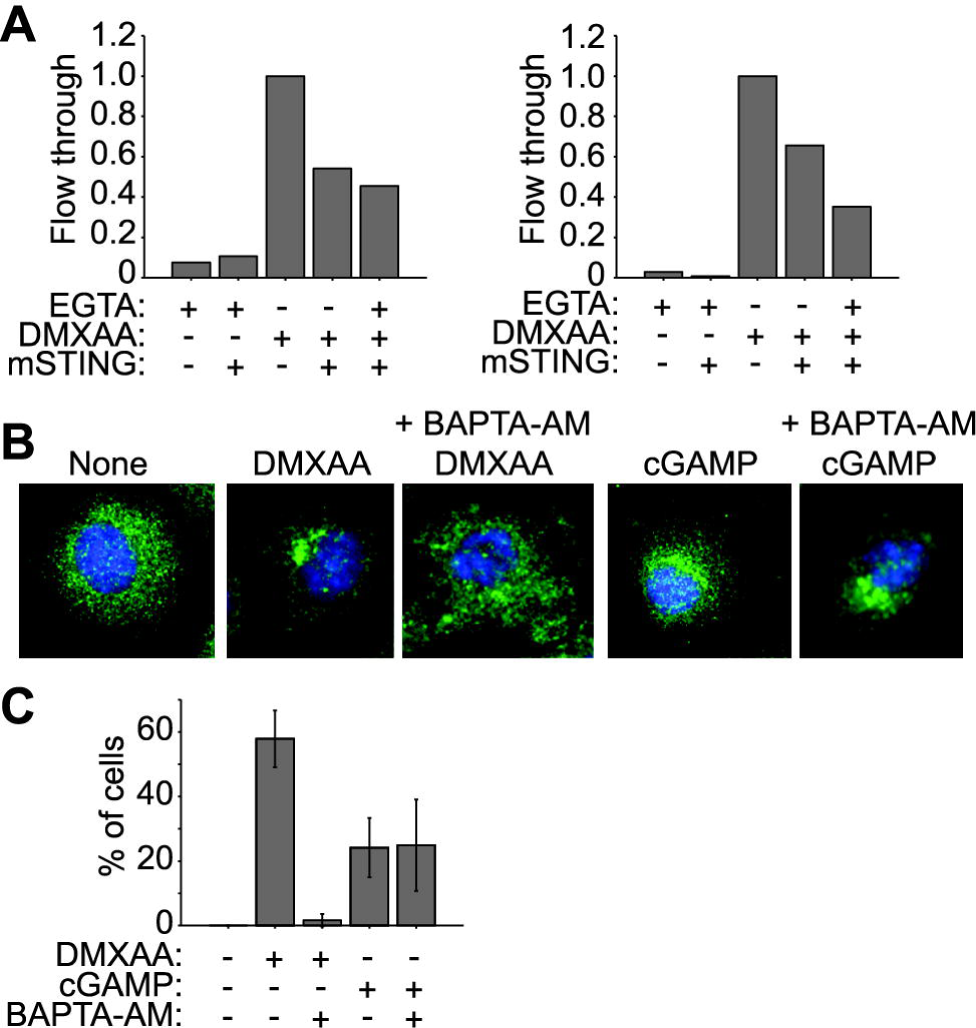
Calcium chelation does not interfere with STING (139-378) and DMXAA or cGAMP interaction *in vitro.* (A) Centrifugal-filtration assay. Mouse STING (139-378) was mixed with small molecules as indicated and loaded onto a 3kD cutoff filter. After centrifugation, absorbance of the flow-through was measured at 245nm for DMXAA and 254nm for cGAMP. (B) Cells were pretreated with BAPTA-AM (30μM) for 30 min and treated with DMXAA (250μM) or cGAMP (50μM) as indicated for an hour. Cells were subjected to paraformaldehyde fixation for immuno-staining. Green represents STING and blue (DAPI) represents the nucleus. (C) Quantification of the number of cells that showed STING re-localization in each condition.

As a further test of calcium involvement in signaling, we measured effects of BAPTA-AM on ligand-induced re-localization of STING. The Barber group showed that STING activation caused it to translocate from the ER to a peri-nuclear compartment [3]. L929 cells were treated with either DMXAA or extracellular cGAMP for one hour and fixed for immuno-fluorescence labeling (Fig. 3B). Both DMXAA and cGAMP caused STING to re-localize as reported, though the effect appeared more dramatic for DMXAA. Approximately 60% of DMXAA treated cells, as compared to ~25% of cGAMP treated cells, showed STING re-localization (Fig. 3C). BAPTA-AM pretreatment blocked translocation caused by DMXAA, but had no effect on translocation caused by extracellular cGAMP (Fig. 3C).

### Evidence against involvement of IP3 and IP3 receptors

Increased cytosolic calcium during signaling usually comes from one of two sources, extracellular calcium via membrane calcium channels or calcium in the ER via IP3 receptors. Extracellular EGTA did not block calcium pulses or signaling induced by DMXAA, implicating IP3 receptors. We first measured IP3 levels, to test if they were increased by DMXAA (Fig. 4A). We used a standard commercial assay that measures IP1, which is formed by rapid hydrolysis of IP3, and is stable in the presence of lithium ions. DMXAA did not cause a measureable increase in IP3, unlike the positive control UDP, which acts as a purinergic GPCR (G protein coupled receptor) ligand in this assay (Fig. 4A). We next tested the effect of two standard IP3 receptor inhibitors, 2-aminoethoxydiphenylborane (2-APB) and Xestospongin C (Xest.C). Neither inhibited DMXAA-induced TBK1-IRF3 activation, while BAPTA-AM did (Fig. 4B). These results suggest that IP3 and IP3 receptors are not involved in generating the calcium transients induced by DMXAA.

**Figure 4.**
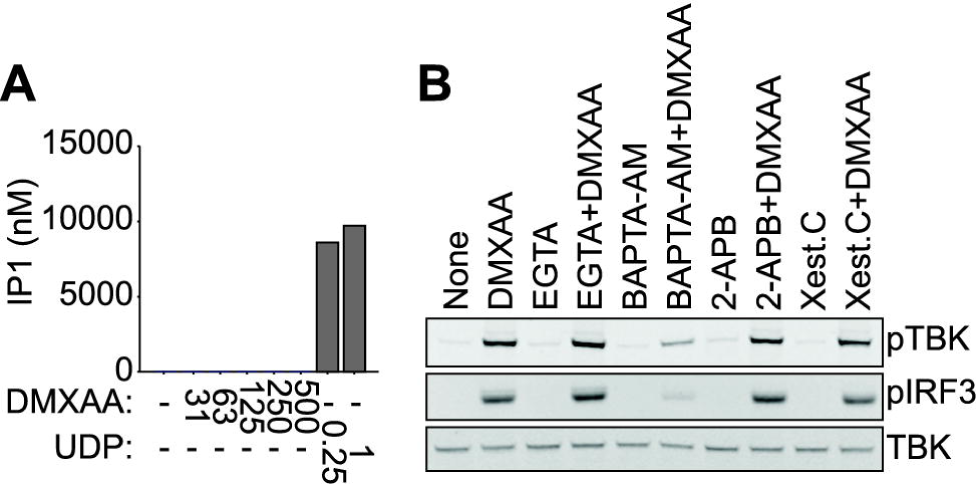
Calcium release through the IP3 receptor was irrelevant for DMXAA induced signal. (A) IP1 assay. Cells were treated with DMXAA or UDP for an hour, lysed, and analyzed with the IPOne assay. (B) Cells were pretreated with EGTA (2mM), BAPTA-AM (30μM), 2-APB (50μM) or Xest.C (10μM) for 30 min and then treated with DMXAA (250μM) for one hour. Phospho-TBK and phospho-IRF3 were measured by western blot.

### Discussion

STING, and ER protein, was discovered as a cyclic dinucleotide receptor that induces type 1 interferon gene expression [27]. We wondered why it localized to the ER, and hypothesized it might mediate calcium release into the cytosol, since calcium is involved in many other cases of activating immune cells. To test this hypothesis, we measured changes in intracellular calcium levels in response to two STING agonists in two cell lines. Both DMXAA and cGAMP rapidly induced calcium transients in the cytoplasm (Fig. 1). In macrophage-like Raw264.7 cells, calcium levels tended to pulse even in untreated cells, whereas DMXAA promoted a large calcium pulse followed by multiple smaller pulses (Fig. 1C). Our imaging was not sophisticated enough to quantify calcium concentrations or precise temporal control. We took images every 20 to 30 seconds, and would have missed sharp spikes. In more fibroblast-like L929 cells, calcium levels were more stable in untreated cells, and both STING ligands induced a single pulse of cytosolic calcium that was stronger for extracellular cGAMP than DMXAA. This pulse was due to specific actions of the ligands since the inactive analog XAA8Me did not induce pulses. Partial STING depletion (~50%) partially inhibited the calcium pulse in response to DMXAA (~15% reduction) and cGAMP (~36% reduction).

Cytosolic calcium was clearly required for STING signaling induced by the synthetic agonist DMXAA. The intracellular calcium chelator BAPTA-AM, which inhibited calcium increase (Fig. 2A, 2B) also inhibited STING pathway activation (Fig. 2C). However, BAPTA-AM did not prevent activation of the STING pathway by extracellular cGAMP (Fig 2D). Why, then, might calcium be required for STING signaling downstream of DMXAA? One possibility is that calcium is required for STING relocalization (Fig. 3B), and relocalization is required for signaling. Another is involvement of some calcium-dependent kinase or phosphatase.

Extracellular cGAMP mobilized calcium more effectively than DMXAA, but BAPTA-AM did not inhibit signaling by this ligand. The reasons for this difference are unclear. Perhaps the two agonists induce different conformational effects in STING, so that DMXAA induces a downstream signal that depends on calcium, while cGAMP induces one that does not, because, for example, the signal strength is higher. However, we see equal degrees of downstream pathway activation for both ligands when they are added at saturating concentration (Fig. 2C, 2D). An alternative hypothesis is that DMXAA is a pure STING ligand, and in this case STING signaling is completely calcium dependent. Extracellular cGAMP, in contrast, might bind to other receptors that transduce a signal to TBK1 and IRF3 by a calcium-independent mechanism, for example using elements of the inflammasome pathway [17].

Finally, what are the likely mechanisms by which calcium enters the cytoplasm following DMXAA treatment, which does appear to act mostly via STING? Neither IP3 nor the IP3 receptor is obviously involved (Fig. 4). We propose that STING itself may act as a ligand gated calcium channel in the ER, or that it interacts with some other ER channel. This hypothesis could explain why STING resides in the ER as a multi-pass transmembrane protein, and seems worthy of further testing.

## Acknowledgments

We thank Yan Feng, John Tallarico at Novartis Institutes for Biomedical Research for providing the flavonoids, Kristin Krukenberg for helpful discussion and critical reading of the manuscript. We also thank the Nikon Imaging Center at Harvard Medical School for help on all microscopy experiments.

